# Negative Overgeneralization is Associated with Anxiety and Mechanisms of Pattern Completion in Peripubertal Youth

**DOI:** 10.1101/2020.01.27.921742

**Authors:** Dana L. McMakin, Adam Kimbler, Nicholas J. Tustison, Jeremy W. Pettit, Aaron T. Mattfeld

**Affiliations:** Cognitive Neuroscience Program, Department of Psychology, Florida International University, Miami, FL 33199 USA; Clinical Science Program, Department of Psychology, Center for Children and Families, Florida International University, Miami, FL 33199 USA; Department of Radiology and Medical Imaging, University of Virginia, Charlottesville, VA 22903 USA; Center for Children and Families, Florida International University, Miami, FL 33199 USA

**Keywords:** anxiety, adolescence, generalization, hippocampus, mPFC, fMRI

## Abstract

**BACKGROUND:** This study examines neural mechanisms of negative overgeneralization in peri-puberty to identify potential contributors to escalating anxiety during this sensitive period. Theories suggest that weak *pattern separation* (a neurocomputational process by which overlapping representations are made distinct, indexed by DG/CA3 hippocampal subfields) is a major contributor to negative overgeneralization. We alternatively propose that neuromaturation related to generalization and anxiety-related pathology in peri-puberty predicts contributions from strong *pattern completion* (a partial match of cues reinstates stored representations, indexed by CA1) and related modulatory mechanisms (amygdala, medial prefrontal cortices [mPFC]).

**METHODS:** Youth (N=34, 9-14 years) recruited from community and clinic settings participated in an emotional mnemonic similarity task while undergoing MRI. At Study, participants indicated the valence of images; at Test, participants made an ‘old/new’ recognition memory judgment. Critical lure stimuli, that were similar but not the same as images from Study, were presented at Test, and errors (“false alarms”) to negative relative to neutral stimuli reflected negative overgeneralization. Univariate, multivariate, and functional connectivity analyses were performed to evaluate mechanisms of negative overgeneralization.

**RESULTS:** Negative overgeneralization was related to greater and more similar patterns of activation in CA1 and both dorsal and ventral mPFC for negative relative to neutral stimuli. At Study, amygdala increased functional coupling with CA1 and dorsal mPFC during negative items that were later generalized.

**CONCLUSIONS:** Negative overgeneralization is rooted in amygdala and mPFC modulation at encoding and pattern completion at retrieval. These mechanisms could prove to reflect etiological roots of anxiety that precede symptom escalation across adolescence.

## INTRODUCTION

Negative overgeneralization is a core feature of anxiety when responses to one aversive situation (e.g., severe weather) spread to non-aversive situations that share stimulus features (e.g., breezy day). Negative overgeneralization can theoretically involve either weak *pattern separation* (a neuro-computational process by which overlapping or similar representations are made distinct) leading to poor behavioral discrimination between threat and safety cues/contexts; or strong *pattern completion* (a neuro-computational process by which a partial match of cues can reinstate previously stored representations) leading to overactive behavioral generalization from threat to safety cues and contexts. There has been a strong focus on pattern separation as a key driver of poor discrimination, with some empirical support in adult populations (for review, see 1). However, neuro-maturational changes during the transition from childhood to adolescence (peri-puberty) potentiate neural mechanisms (e.g. hippocampal subregions, amygdala) that normatively favor pattern completion (e.g. 2–4) and therefore could drive negative overgeneralization among vulnerable youth. Ultimately, these mechanisms may prove to reflect etiological roots of anxiety that immediately precede the known escalation of symptoms across adolescence (5).

We draw specifically on three lines of converging evidence to support and test our theory that pattern completion and related hippocampal and modulatory neural mechanisms drive negative overgeneralization in peri-puberty. First, normative developmental changes enhance pattern completion and generalization, relative to separation and discrimination, in peri-puberty. Young children have a strong tendency to generalize information to allow for rapid learning and broad threat and reward detection (6–8). As children transition through adolescence, generalization is increasingly complemented by precision in the ability to discriminate from among similar events (6,9,10). These behavioral changes coincide with neuro-maturational changes in hippocampal subfields that contribute to pattern completion and separation. The Cornu Ammonis (CA)3 subfield heavily supports pattern completion, while the dentate gyrus (DG) supports pattern separation (11,12). In human fMRI studies, activations in CA3/DG subregions are combined to reflect pattern separation; while activations in CA1, which receives input from CA3, reflect pattern completion (13). The CA1 develops earlier while the DG/CA3 exhibits a protracted developmental trajectory through early adulthood (2) and recent studies suggest that changes in DG/CA3 volume are associated with the developmental shift from generalization to discrimination from late childhood through early adulthood (2–4). However, whether or not a relative strength in pattern completion at the neuromaturational inflection point of peri-puberty contributes to overgeneralization in youth with, or at risk for, anxiety is unknown.

Second, neural regions that modulate hippocampal function (e.g. amygdala, mPFC) also evidence protracted development in ways that may further tip systems toward pattern completion of emotional stimuli. Amygdala-based emotional arousal has been shown to enhance the creation of long-lasting memories (14), and increases in generalization are reliably identified following negative (15), aversive (16), or threatening (17) events. Moreover, lesions of the amygdala result in selective deficits of gist but not detailed memories (18). Protracted maturation of the amygdala and related modulatory circuitry leads to greater reactivity of amygdala in peri-puberty, relative to mid-to late-adolescence and adulthood (19), which could amplify tendencies toward generalization. The mPFC also undergoes protracted development, and exercises control over the specificity of hippocampal memories (20) and can both potentiate (dorsal mPFC; dmPFC) and inhibit (ventral mPFC; vmPFC) fear memory (21,22)—both have been implicated in negative generalization during fear conditioning (e.g., 23–31). Taken together, these maturational changes could lead to a mnemonic system that is readily influenced by emotional arousal, and shows a tendency toward generalization.

Finally, youth with anxiety evidence differences in generalization and related neural mechanisms in ways that could lead to a maturational inflection point and contribute to worsening symptoms. For example, a recent study demonstrated a wider generalization gradient for anxious youth in response to a conditioning task where a loss or gain was associated with well separated tones and a tone probe was subsequently used to assess generalization curves (32). Moreover, this study demonstrated normative elevation in generalization among the child group, relative to adolescents, but in youth with anxiety the pattern was reversed. This reversal may suggest a neuro-maturational inflection point for negative overgeneralization that uniquely affects vulnerable youth.

Although there are no available neuroimaging studies of negative overgeneralization in youth, it is known that youth with anxiety show increased activation of the amygdala and reduced connectivity with vmPFC in response to threatening or ambiguously threatening stimuli (33) which could amplify maturational tendencies toward pattern completion and negative overgeneralization during this sensitive period.

To examine neural mechanisms of negative overgeneralization at this developmental cusp of escalating risk, we sample peri-pubertal youth (9-14 years) across a continuum of anxiety symptoms to maximize variability in negative overgeneralization. We use a declarative memory paradigm with high resolution fMRI that is well established for distinguishing behavioral generalization and discrimination, and for visualizing hippocampal subfields associated with pattern completion and separation. We hypothesize that negative overgeneralization will be associated with neural mechanisms of pattern completion (greater CA1 activation) and modulatory effects of amygdala and mPFC, and that anxiety severity will be associated with greater negative overgeneralization in terms of behavior and activation in proposed neural mechanisms.

## METHODS AND MATERIALS

### Participants

Volunteers, ages 9-14 years, were recruited from anxiety clinic referrals and the community and paid for their participation. The local IRB approved study procedures and participants completed informed consent and assent. Participants were screened for major medical and psychiatric exclusionary comorbidities (e.g., current depressive episode, bipolar disorder, post-traumatic stress disorder, attention-deficit/hyperactivity disorder, conduct disorder, oppositional defiant disorder, psychotic disorders, obsessive compulsive disorder). Forty-eight participants met eligibility requirements. All participants were right-handed and had normal or corrected to normal vision. Participants were dropped from final analyses due to errors in data collection at either Study or Test (n = 4), excessive motion (n = 2; >20% of trials were flagged for removal), failed to show up for their scheduled Study or Test session (n = 2), or poor task performance (n = 6; hit rate for targets was 1.5 SD below the average performance), leaving N = 34 (11.4 ± 2.0 years, 16 female) for the final sample. Anxiety severity was assessed using the Pediatric Anxiety Rating Scale (PARS-6; 34).

### General design procedures

The study design included randomization of participants to sleep (n = 16) or wake (n = 18) conditions to evaluate the role of sleep in emotional memory consolidation. In both conditions, Study and Test scans were separated by a 12-hour retention period. The timing for the two conditions differed such that the sleep condition completed Study scan in the evening and Test scan in the morning, while the wake condition completed Study scan in the morning and Test scan in the evening. The effects of sleep versus wake conditions on negative overgeneralization will be detailed in a future report but are outside of the scope of the current study which is instead focused on general mechanisms of negative overgeneralization.

### Emotional Memory Task Procedures

The emotional mnemonic similarity task (35) consisted of two phases: Study and Test (Figure 1A). During Study, participants viewed (2s) emotional and non-emotional scenes in a randomized order. Participants were instructed to indicate the valence (negative, neutral, or positive) of each image with one of three button responses. Each scene was followed by a white fixation cross in the middle of a black screen of variable duration (2-6s) before the onset of the next trial. At Study, participants saw 145 stimuli (48 negative, 47 neutral, 50 positive) equally divided between two scanning runs lasting approximately 8min. Participants returned to the scanner after a 12hr delay for Test, at which point they were given a surprise memory test. Participants were presented with scenes which were viewed at Study (targets), scenes that were similar but not identical to the images presented at Study (lures), and completely new scenes (foils). Each scene was presented in the middle of the screen for 2s, followed by a variable inter-stimulus-interval (2-6s) during which a white fixation cross was presented in the middle of a black screen. Participants were instructed to indicate whether items were “old” or “new” by pressing one of two buttons. The instructions indicated that for an item to be endorsed as “old” it had to be the *exact same* image that was presented at Study. A total of 284 stimuli were presented at Test: 16 each negative, neutral, and positive targets, 32 negative, 33 neutral, and 33 positive lures, and 42 negative, 49 neutral, 48 positive foils.

**Figure 1.**
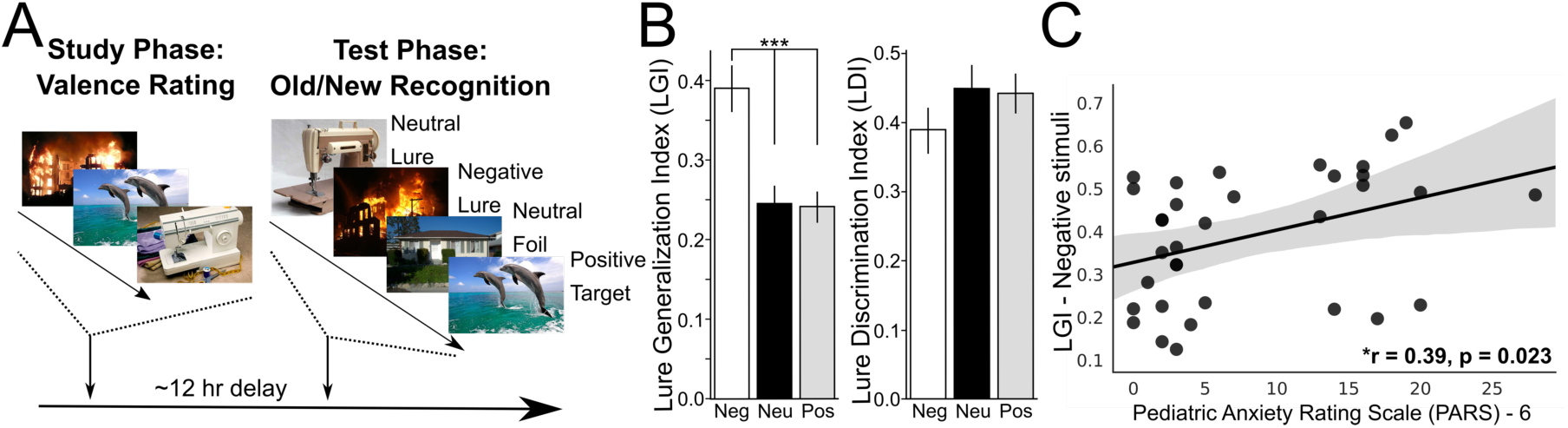
Increased false alarms to negative lures in peripubertal youth. Timeline of experimental procedures and lure behavioral performance. (**A**) The scanning portion of the experiment consisted of two phases: 1) A Study phase during which participants rated the valence of each image. Images were presented for 2s and were separated by a jittered inter-stimulus-interval (ISI: white fixation cross; 2-6s). 2) During the Test phase participants performed an old/new recognition memory task. Target (images from the Study phase), foil (new pictures), and lure (new images that were similar but not exactly the same as images from the Study phase) stimuli were presented for 2s and separated by a jittered ISI (2-6s). (**B**) The Lure Generalization Index (LGI): p(‘Old’|Lure) – p(‘Old’|Foil) – was significantly greater for negative compared to neutral (*t*(33) = 5.07, *P* = 1.4×10^−5^) and positive (*t*(33) = 6.44, *P* = 2.6×10^−7^) stimuli. A similar difference was not observed (all *P* > 0.10) for the Lure Discrimination Index (LDI): p(‘New’|Lure) – p(‘New’|Target). Data are represented as a mean ± SEM. (**C**) Negative generalization (Negative LGI) was positively associated with anxiety severity (PARS-6) (*r* = 0.39, *P* = 0.023). Neg = Negative; Neu = Neutral; Pos = Positive; *** *P* < 0.0001.

### Neuroimaging Data Collection and Preprocessing

Neuroimaging data were collected on a 3T Siemens MAGNETOM Prisma scanner with a 32-channel head coil at the Center for Imaging Science at Florida International University. A T2*-weighted sequence (TR = 993ms, TE = 30ms, flip angle = 52°, FOV = 216mm, 56 axial slices, slice acceleration = 4, voxel size = 2.4mm isotropic) in addition to a T1-weighted sequence (TR = 2500ms, TE = 2.9ms, flip angle = 8°, FOV = 256mm, 176 sagittal slices, voxel size = 1mm isotropic) were collected. During each run of the emotional memory task acquisition began after the first four volumes were collected. A total of 448 whole brain volumes were acquired at Study and 454 volumes were collected at Test. Neuroimaging preprocessing procedures, including motion correction, co-registration, outlier detection, spatial filtering, and template normalization, were utilized (for detailed description see Supplemental Information).

### Behavioral Data Analysis

To assess generalization, we calculated how likely participants were to false alarm to lure items correcting for a tendency to call new stimuli ‘old,’ p(‘Old’|Lure) – p(‘Old’|Foil), called the lure generalization index (LGI). To evaluate discrimination, we computed the likelihood that participants correctly rejected the same stimuli corrected by their propensity to endorse target stimuli ‘new’, p(‘New’|Lure) – p(‘New’|Target), and referred to this measure as the lure discrimination index (LDI). Similar procedures have been used in prior work (36–38). Pairwise comparisons were performed across the different stimulus valences for both the LGI and LDI scores. Bonferroni corrections were used to correct for inflated Type I error across pairwise comparisons.

### MRI Data Analysis

#### Anatomical Regions of Interest (ROIs)

The amygdala was defined by binarizing the subcortical FreeSurfer segmentation, while the dmPFC and vmPFC were defined by binarizing the following cortical segmentations – dmPFC: superior frontal, caudal anterior cingulate, and rostral anterior cingulate; vmPFC: medial orbitofrontal.

Hippocampal subfield segmentation was performed using a consensus labeling approach based on an in-house atlas set of 19 T1 MPRAGE scans (0.75mm isotropic) and their corresponding T2-FSE scans, acquired in an oblique orientation perpendicular to the long axis of the hippocampus (0.47×0.47mm^2^ in-plane, 2.0mm slice thickness). Expert manual segmentations were applied to this atlas set based on highly reliable and published protocols (39,40) and comprised the following bilateral ROIs: DG/CA3, CA1, and subiculum. Details can be found in Supplemental Information.

All masks were back-projected to functional space using the inverse of the co-registration transformation for data analyses.

### Neuroimaging Data Analysis

Functional neuroimaging data were analyzed using a general linear model approach in FSL. Separate models were created for the Study and Test data to evaluate the neurobiological correlates of negative generalization. Both the Study and Test models included the following regressors of no interest: motion (x, y, z translations; pitch, roll, yaw rotation), the first and second derivatives of the motion parameters, normalized motion, first through third order Lagrange polynomials to account for low frequency changes in the signal, as well as a regressor for each outlier time-point that exceeded outlier thresholds. The Test model included 12 regressors of interest: lure false alarms (FAs), lure correct rejections (CRs), target hits, and target misses. All regressors were separately modeled for each valence (negative, neutral and positive). The Study model included a similar combination of regressors, however, these were scenes that would subsequently be target hits and misses or be replaced by similar lures and subsequently correctly rejected or false alarmed. To investigate signals related to negative generalization we focused analyses on negative compared to neutral lure trials that were (subsequently) false alarmed. Correlation analyses were performed using a beta-series approach (41) to assess changes in task-based functional connectivity. Lastly, a multivariate approach, representational similarity analysis (RSA), was utilized to evaluate the relation between behavioral generalization and neural measures of pattern similarity. See Supplementary Information for a detailed description of correlational and multivariate analyses.

## RESULTS

### Increased false alarms to negative lures in peripubertal youth

To examine negative overgeneralization behaviorally, we compared LGI between negative, neutral, and positive stimuli. Generalization was significantly greater for negative compared to both neutral (*t*(33) = 5.07, *P* = 1.4×10^−5^) and positive (t(33) = 6.44, *P* = 2.6×10^−7^) stimuli, which were not significantly different from one another (*t*(33) = −.16, *P* = .87) (Figure 1B). The same comparisons were performed using LDI. No significant differences in discrimination were evident when comparing the different stimulus valences (LDI_Neg v. Neu_: *t*(33) = −1.56, *P* = .12; LDI_Neg v. Pos_: *t*(33) = −1.66, *P* = 0.106; LDI_Pos v. Neu_: *t*(33) = −.19, *P* = 0.85). To evaluate the relationship between negative overgeneralization and anxiety we correlated Negative LGI with PARS-6. Participants with greater anxiety severity were more likely to generalize negative information (*r* = 0.39, *P* = 0.023) (Figure 1C).

### MRI Results

#### Negative generalization is associated with greater activation in the mPFC and CA1

To investigate neurobiological mechanisms supporting negative overgeneralization, we examined differences in activations for negative relative to neutral lure stimuli that were incorrectly identified as old in anatomically defined ROIs. We found greater activation in the CA1 for negative compared to neutral lures (*t*(33) = 3.51 *P* = 0.001) (Figure 2A). Activations between negative and neutral lures in the DG/CA3 did not significantly differ following corrections for multiple comparisons but exhibited a trend towards greater activation for negative relative to neutral items (*t*(33) = 2.29, *P* = 0.03), in the opposite direction predicted based on a failure to pattern separate (Figure 2B). Both the dmPFC (*t*(33) = 2.74, *P* = 0.009) and vmPFC (*t*(33) = 3.24, *P* = 0.002) exhibited greater activation for negative when compared to neutral lures (Figure 2C,D).

**Figure 2.**
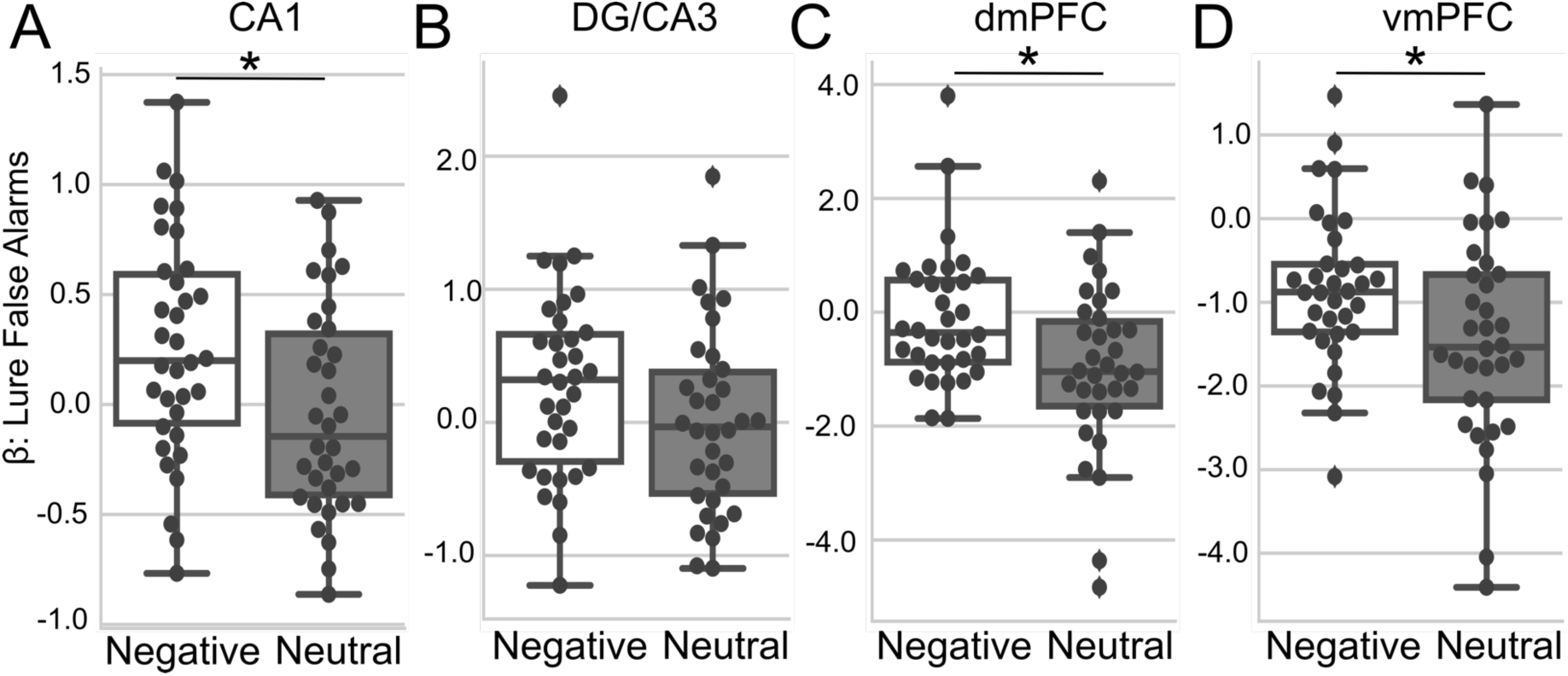
Negative generalization is associated with greater activation in the CA1 and mPFC. Anatomical region-of-interest analysis showing hippocampal subfield (CA1 and DG/CA3) and both dorsal medial prefrontal cortex (dmPFC) and ventral medial prefrontal cortex (vmPFC) activations. (**A**) The CA1 hippocampal subfield (*P* = 0.001) exhibited greater activations for negative lure false alarms relative to neutral lure false alarms. (**B**) The DG/CA3 subfields (*P* = 0.03) exhibited a trend towards greater activations for negative lure false alarms relative to neutral lure false alarms. (**C**,**D**) Both the dmPFC (*P* = 0.009) and the vmPFC (*P* = 0.002) showed greater activations for negative compared to neutral lure false alarms. Boxplots reflect median and interquartile range. Error bars reflect maximum and minimum range. *Significant following Bonferroni correction for multiple comparisons.

To further evaluate whether negative overgeneralization was a result of impaired discrimination resulting from poor pattern separation we compared activations in the same regions for negative and neutral lure stimuli that were identified as ‘New’ (e.g., CR). Amongst ROIs no regions exhibited a significant difference between negative and neutral lures following corrections for multiple comparisons (CA1: *t*(33) = 2.32, *P* = 0.03; DG/CA3: *t*(33) = 2.50, *P* = 0.02; vmPFC: *t*(33) = 0.29, *P* = 0.76; dmPFC: *t*(33) = 0.39, *P* = 0.69) (Supplemental Figure 1). In fact, similar to FAs in the DG/CA3, there was a trend towards greater activation in the CA1 and DG/CA3 for negative relative to neutral lures – opposite the predicted difference based on an assumption of impaired pattern separation.

### A subcortical and cortical network contribute to negative overgeneralization

To evaluate the contribution of other brain areas to negative overgeneralization outside our anatomical ROIs, we performed an exploratory whole-brain analysis. We compared negative to neutral lure FAs. Similar to our anatomical ROI approach, clusters in the hippocampus survived corrections for multiple comparisons. We also observed clusters in the bilateral amygdala and midline thalamus. Clusters in the bilateral mPFC, posterior cingulate cortex extending into the precuneus, inferior parietal lobule, and middle temporal gyrus were observed (Figure 3).

**Figure 3.**
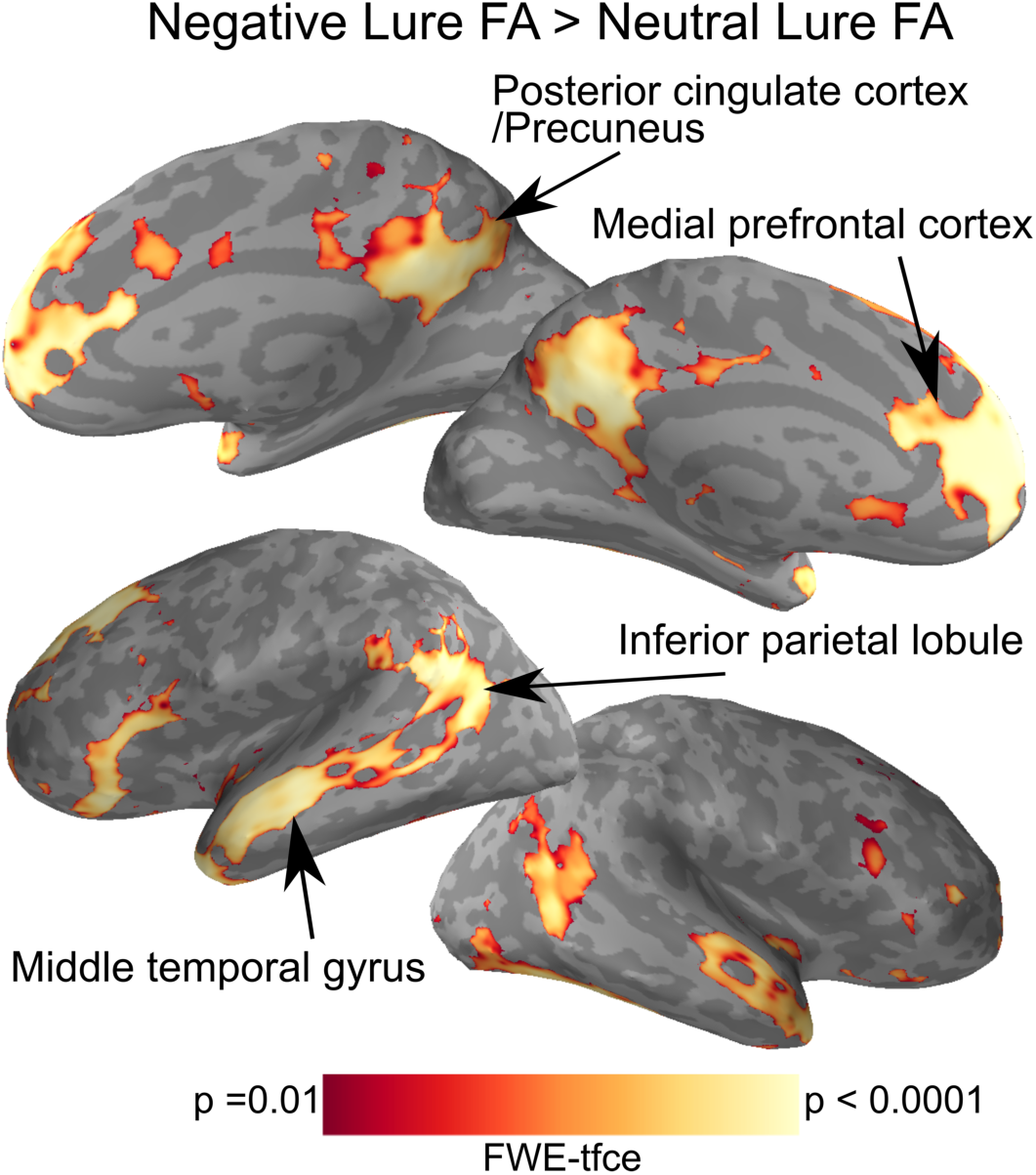
A subcortical and cortical network contribute to negative overgeneralization. Whole brain exploratory analysis showing regions that exhibited greater activations for negative relative to neutral lure false alarms (FA) in the posterior cingulate cortex extending into the precuneus, medial prefrontal cortex, inferior parietal lobule, and middle temporal gyrus. FWE-tfce corrected *P* < 0.01.

### Enhanced functional coupling with amygdala during encoding

We next assessed the functional correlations between amygdala and target regions at Study. We observed enhanced functional coupling between the amygdala and both the CA1 (amygdala ↔ CA1: *t*(33) = 2.9, *P* = 0.007) and the dmPFC (amygdala ↔ dmPFC: *t*(33) = 2.6, *P* = 0.013) for negative stimuli compared to neutral stimuli that were subsequently FA but only a trend for the amygdala and the vmPFC (amygdala ↔ vmPFC: *t*(33) = 1.6, *P* = 0.13) (Figure 4). No significant difference in functional connectivity between the same regions was observed when comparing negative and neutral stimuli that were subsequently CR (amygdala ↔ CA1: *t*(33) = 1.03, *P* = 0.32; amygdala ↔ dmPFC: *t*(33) = −0.46, *P* = 0.64; amygdala ↔ vmPFC: *t*(33) = −0.76, *P* = 0.45) (Supplemental Figure 2).

**Figure 4.**
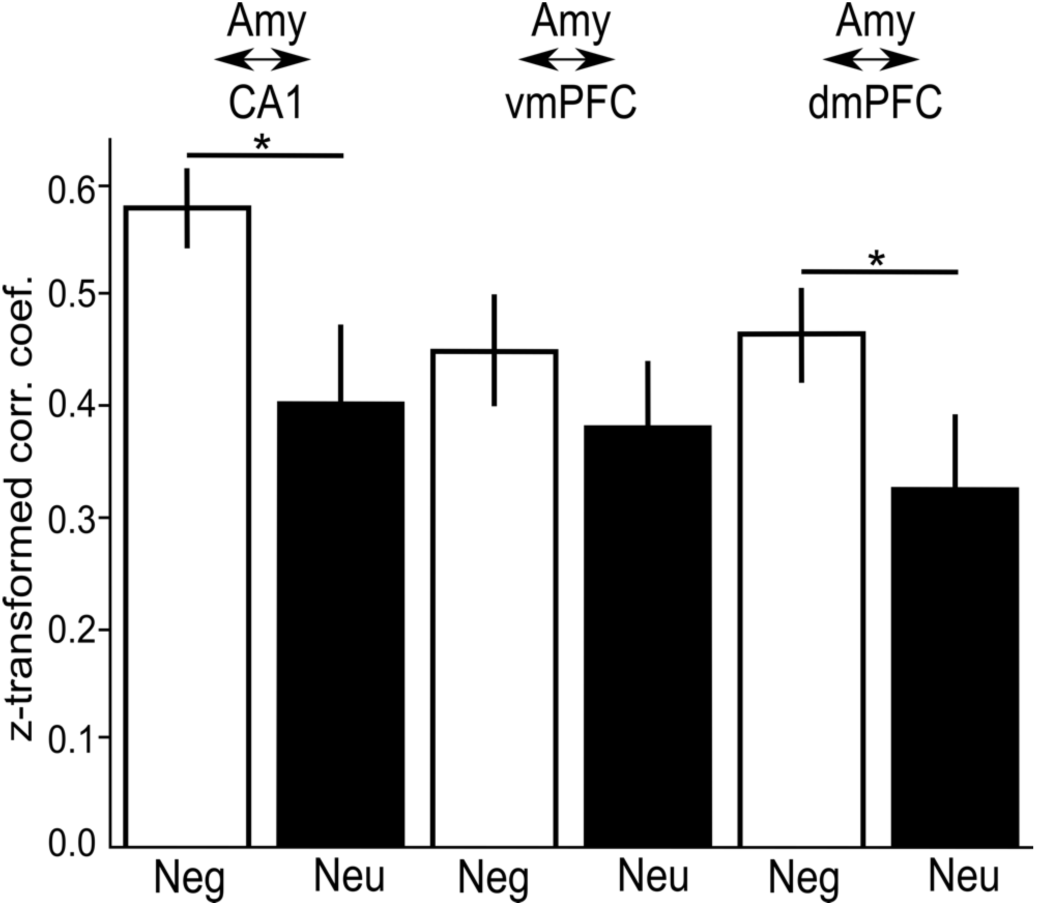
Enhanced functional coupling with amygdala during Study. Task based functional connectivity during the Study phase between the amygdala (Amy) and three target regions: CA1, ventral medial prefrontal cortex (vmPFC), and dorsal medial prefrontal cortex (dmPFC) for items that were subsequently replaced by lures and false alarmed. Increased functional connectivity was observed between the Amy ↔ CA1 (*P* = 0.007) and the Amy ↔ dmPFC (*P* = 0.013) for negative (Neg) relative to neutral (Neu) stimuli that were replaced by similar lure stimuli and subsequently false alarmed. Data are represented as a mean ± SEM. *Significant following Bonferroni correction for multiple comparisons.

### Increased representational similarity in the mPFC and left CA1 associated with enhanced negative overgeneralization

To evaluate the relationship between patterns of brain activation and our behavioral indices of generalization (i.e., LGI) we utilized RSA to examine the correlation between target hit and lure false alarm similarity and LGI scores. Negative LGI increased as both vmPFC (*r* = 0.53, *P* = 0.001) and dmPFC (*r* = 0.44, *P* = 0.009) similarity between negative target hits and lure false alarms increased (i.e., became more similar) (Figure 5A,B). A similar pattern was observed in the left CA1 (*r* = 0.46, *P* = 0.006) (Figure 5C) – negative LGI increased as similarity in the left CA1 increased – but not in the right CA1 (*r* = 0.08, *P* = 0.65) (Figure 5D).

**Figure 5.**
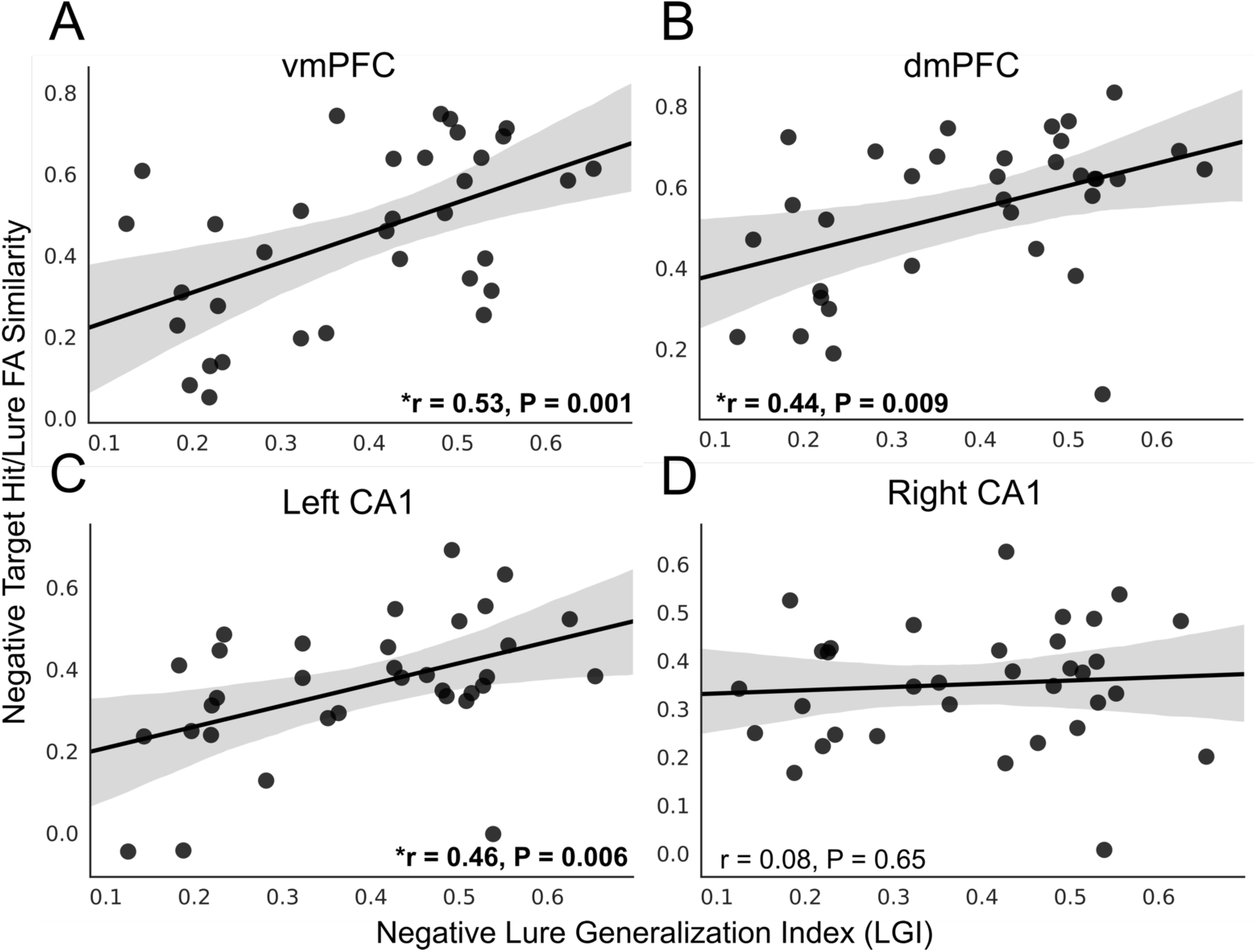
Increased representational similarity in the mPFC and left CA1 associated with enhanced negative overgeneralization. Correlations between negative target hit and lure false alarm (FA) similarity in the ventral medial prefrontal cortex (vmPFC), dorsal medial prefrontal cortex (dmPFC), and bilateral CA1 and LGI behavioral performance. (**A**,**B**) Both the vmPFC (*P* = 0.001) and the dmPFC (*P* = 0.008) exhibited a significant correlation between negative target hit/lure FA similarity and negative LGI performance. As the similarity between target hit and lure FA activations increased (i.e., became more similar) negative LGI performance increased. (**C**,**D**) A similar correlation was identified in the left CA1 (*P* = 0.006) but not the right CA1 (*P* = 0.65). *Significant following Bonferroni correction for multiple comparisons.

Both lure FAs and target hits were endorsed with an ‘old’ response, thus the similarity in activations may be related to the same responses across the two conditions. To evaluate this confound, we performed the same analyses using positive and neutral stimuli. No significant relationship between pattern similarity and behavior, despite the same response type, was observed for positive stimuli (left CA1: *r* = −0.015, *P* = 0.93; right CA1: *r* = −0.16, *P* = 0.35; vmPFC: *r* = 0.09, *P* = 0.59; dmPFC: *r* = 0.19, *P* = 0.27) (Supplemental Figure 3) and only a nonsignificant trend for neutral stimuli was observed (left CA1: *r* = 0.40, *P* = 0.018; right CA1: *r* = 0.30, *P* = 0.08; vmPFC: *r* = 0.29, *P* = 0.09; dmPFC: *r* = 0.13, *P* = 0.46; Supplemental Figure 4).

### Relationship between anxiety severity and brain measures

Lastly, we explored the relationship between the identified neurobiological mechanisms of negative overgeneralization and anxiety symptom severity. No significant relationship between anxious symptoms (e.g., PARS-6) and univariate activations amongst our ROIs was evident (CA1: *r* = 0.04, *P* = 0.84; vmPFC: *r* = 0.18, *P* = 0.32; dmPFC: *r* = −0.03, *P* = 0.87). Functional connectivity during the Study phase between the amygdala ↔ CA1 and the amygdala ↔ dmPFC for negative items that were subsequently false alarmed exhibited a trend, whereby greater connectivity was associated with more severe symptoms of anxiety (*r* _AMY↔CA1_ = 0.29, *P* = 0.09; *r* _AMY↔dmPFC_ = 0.35, *P* = 0.04) (Supplemental Figure 5A). A similar trend in our multivariate measure was noted, wherein representational similarity in the left CA1 between negative target hits and negative lure false alarms was elevated for participants with greater levels of anxiety (*r* = 0.32, *P* = 0.06) (Supplemental Figure 5B). This relationship was not evident in the vmPFC (*r* = 0.02, *P* = 0.85) or the dmPFC (*r* = 0.44, *P* = 0.80).

## DISCUSSION

Behavioral and neurobiological data from an emotional mnemonic similarity task converged to support the notion that negative overgeneralization during peri-puberty results from enhanced pattern completion. Specifically, results indicated greater behavioral generalization (e.g., LGI) and neural activation in CA1 and mPFC when generalizing negative relative to neutral lures. Neurobiological mechanisms of negative overgeneralization are modulated by interactions between the amygdala with target regions during encoding, as evidenced by greater amygdala ↔ CA1 and dmPFC connectivity during the Study phase for negative (relative to neutral) stimuli that were subsequently generalized. Representational similarity analyses also showed that more similar neural activation patterns in left CA1 and mPFC were associated with greater behavioral generalization for negative stimuli. Greater negative overgeneralization was associated with elevated anxiety, however, only a trend with neurobiological mechanisms was identified. Together, these results outline candidate neurobiological mechanisms underlying negative overgeneralization. Contributions from these neural circuits and increasing symptoms of anxiety during peri-puberty highlight the potential susceptibility and plasticity of these mechanisms during this developmental window, providing a substrate from which anxiety and related disorders could emerge or escalate.

Activations in *a priori* ROIs were associated with the computational process of pattern completion. Prior studies have identified the hippocampus as an important contributor to negative overgeneralization (23,24,29–31,42), however the contributions of hippocampal subfields have remained underspecified. Using high resolution imaging, we observed greater activation in the CA1 for negative versus neutral lure false alarms – consistent with generalization through pattern completion. The ‘generalization’ interpretation of CA1 activation in our study is in line with computational models that have proposed CA1 activity may generalize following partial or incomplete pattern completion in the CA3 (43,44). In contrast, the DG/CA3, a subfield of the hippocampus that contributes to successful discrimination through pattern separation (11,12), exhibited a trend towards greater activation for negative compared to neutral lure false alarms and correct rejections – a directionality that is inconsistent with predictions based on impairments in pattern separation.

Theoretical models have emphasized poor pattern separation and related deficits in behavioral discrimination in models of aging as well as both typical and clinical adult populations (1). Our results challenge the transcendence of pattern separation, supporting the notion that neural mechanisms underlying the tendency to generalize or the failure to discriminate vary across developmental periods. The divergent mechanisms across development may be further complicated by pathology. For example, negative generalization is exacerbated in adolescents compared to children but only in those with a diagnosis of anxiety (45).

We also found support for involvement of mPFC in negative overgeneralization. However, both the dmPFC, as well as the vmPFC exhibited greater activation for negative compared to neutral lure false alarms suggesting that the functional distinction between these regions identified during aversive conditioning paradigms (21,22,25,29,30,42,46) may not hold for more complex tasks and/or this developmental period. Similar conclusions have been drawn in rodent models using context fear conditioning paradigms (21,23,24,42,46). Negative overgeneralization was not limited to the CA1 and mPFC, rather a broad network including regions like the amygdala, posterior cingulate cortex/precuneus, and midline thalamus also exhibited greater activations for negative relative to neutral lure stimuli that were false alarmed lending support to the hypothesis that a broad network governs negative overgeneralization.

Taken together, these findings stand in contrast to existing literature pointing to failures in pattern separation as a neurocomputational driver of negative overgeneralization. However, prior studies have been conducted primarily with rodent models, healthy adults, aged adults, or adults with subclinical depressive symptoms (see 1 for a review). Our focus on peri-pubertal youth and sampled across a full range of anxious symptoms to maximize variability in the construct of interest (i.e. negative overgeneralization), likely account for these differences.

During the Study phase of our experiment we observed increased functional correlations between the amygdala and both the CA1 and dmPFC for negative relative to neutral items that were subsequently replaced by a lure and false alarmed. These regions are important for generalization and fear potentiation respectively (22,47). Greater functional coupling for similar stimuli that were correctly rejected was not observed, discounting the possibility that the amygdala is more correlated with negative stimuli regardless of behavioral performance. These results support our theory that the amygdala at encoding biases neurobiological mechanisms towards generalization, consistent with prior studies of emotions preferentially enhancing gist rather than detailed memories (48).

As a complementary analytic approach, we used representational similarity analyses to investigate the contributions of the CA1 and mPFC. We found support for our hypothesis that negative overgeneralization is strongest when the representations of novel lure stimuli resemble the representations of previously encountered items. Consistent with this prediction, negative LGI was greater among individuals with highly similar patterns of activation between negative target hits and lure false alarms. These results are important given that both the hippocampus and mPFC play a role in memory integration and concept formation (49–52). Thus, the relation between the convergent patterns of brain activation and negative overgeneralization behaviorally may reflect mechanisms of pattern completion and memory integration, respectively.

Finally, our exploratory analyses of associations between neurobiological mechanisms of negative overgeneralization and anxiety severity evidenced either weak or absent effects. We believe there could be at least two explanations. First, anxiety disorders are highly heterogeneous, and although negative overgeneralization is one common symptom dimension, it is not directly assessed in the anxiety severity interview (PARS-6). Therefore, the summary score of anxiety severity may be diffuse, contributing to weak associations with neurobiological mechanisms of negative overgeneralization. Additionally, the sample size is likely underpowered to support multiple interactions across response systems. Future work could use measures specifically designed to assess the symptom dimension of negative overgeneralization in a larger sample in order to clarify symptom-mechanism associations.

These data support a neurobiological mechanism underlying negative overgeneralization. Interactions between the amygdala and target regions during initial encoding contribute to the formation of gist like representations. At retrieval, enhanced univariate as well as convergence of multivariate patterns in activation in both the mPFC and CA1 reflect enhanced pattern completion. In this way interactions between the amygdala and mPFC and CA1 at encoding determine their contribution to negative overgeneralization at retrieval. These findings emphasize the importance of considering developmental stages when defining mechanisms of negative overgeneralization, and can further detail working models of anxiety pathogenesis during the transition from childhood to adolescence. Findings such as these also carry implications for novel treatment development. For example, strategies that aim to improve the balance of pattern completion relative to separation processes in relation to emotional experiences in vulnerable youth could help to mitigate overgeneralization at this key maturational inflection point. Such strategies may include those that aim to reduce emotional reactivity and associated amygdala response to aversive experiences (i.e. in the context of exposure therapy) as a means to mitigate modulatory influences on pattern completion and generalization, and/or strategies such as memory specificity training (thus far evaluated mostly in relation to adult depression; e.g. 53) to reduce strong completion tendencies and related overgeneralization of past negative memories. The neural mechanisms of negative overgeneralization in peripubertal youth that were identified in this study provide defined mechanistic targets for developing and refining such treatment strategies from an experimental therapeutics framework.

## ACKNOWLEDGEMENTS

We would like to thank the team at the Center for Imaging Sciences at Florida International University, the EMU project staff, and participant families, for their assistance in collecting these data. We thank Drs. Timothy Allen and Michael Yassa for their useful feedback on the project and manuscript. The research was funded in part by an R01 from the NIMH to DLM and ATM (MH116005).

## DISCLOSURES

The authors declare no competing interests.

## SUPPLEMENTAL INFORMATION

### MRI Preprocessing

The following software packages were utilized for neuroimaging data preprocessing using a custom scripted Neuroimaging in Python (Nipype version 0.12.1; 1) pipeline: Analysis of Functional Neuroimages (AFNI version 16.3.18; 2), FMRIB Software Library (FSL version 5.0.10; 3), FreeSurfer (version 6.0.0; 4), and Advanced Normalization Tools (ANTs version 2.1.0; 5). Cortical surface reconstruction and cortical/subcortical segmentation was obtained for each T1-weighted MPRAGE. Surfaces were inspected for errors, manually edited (if necessary), and resubmitted for reconstruction. Functional volumes that exceeded three standard deviations of the mean intensity within a run were removed and replaced (‘despiked’). Functional data were then 1) motion corrected, aligning all volumes across each run to the middle volume of the first run; 2) co-registered to the structural scan; 3) motion and intensity outliers (>1 mm frame-wise displacement; >3 SD mean intensity) were identified; and 4) spatially filtered with a 4mm kernel using the *SUSAN* algorithm (6).

Each participant’s T1-weighted structural scan was skull-stripped and registered to an MNI-152 template via a rigid body transformation. We used this pass to reduce normalization errors due to large differences in position across participants and to generate a template close to a commonly used reference. The rigid-body transformed structural scans were then used to create a study-specific template using ANTs (10). Following template generation each participant’s skull-stripped brain in FreeSurfer space was normalized to the study template using ANTs non-linear symmetric diffeomorphic mapping (8). The warps obtained from normalization were used to move contrast parameter estimates following fixed-effects modeling into the study-specific template space for group-level tests.

### Hippocampal Subfield Consensus Labeling Approach

To propagate a weighted consensus labeling from the expertly labeled atlas set to one of the unlabeled T1-weighted images of our study cohort, we spatially normalized the atlas set to the unlabeled subject and applied joint label fusion (7). This consensus-based labeling approach as well as the spatial normalization steps employed the ANTs toolkit (8). First, the intra-subject atlas T1/T2 rigid transforms were calculated. In order to minimize the total number of deformable registrations, a “pseudo-geodesic” approach was used to align the data (9). This required the construction of an optimal T1-weighted template representing the average shape/intensity information of the T1 component of the atlas set (10). The deformable transformations between each T1-weighted image of the study cohort and the T1 atlas template were calculated (9). The transformation between the atlas labels and unlabeled study cohort image was then computed by concatenating the T1 _atlas_ /T2_atlas_ rigid transformation, the T1_atlas_ /T1 template deformable transformation, and the T1 template/and T1_subject_ deformable transforms. Once the atlas set was normalized to the unlabeled subject, the regional labeling was determined using weighted averaging where the weighting takes into account the unique intensity information contributed by each atlas member. This approach which combines label information and intensity information has been used in a number of recent publications to segment hippocampal subfields (10–13).

### Task-Based Functional Connectivity Analysis

We used a beta-series correlation analysis (14) to assess the functional interactions between the amygdala and the CA1, dorsal mPFC, and ventral mPFC during encoding. To isolate our trials of interest (negative and neutral stimuli that were subsequently replaced by lures and either false alarmed or correctly rejected) we used a least-squre single (LSS) approach (15). A general linear model was run for each trial of interest. Other regressors in the models included the typical task-based regressors (minus the relevant trial of interest) and regressors of no interest accounting for motion and low frequency noise. The resulting contrast parameter estimates for each isolated trial of interest were constructed into a beta-series and averaged across voxels within each region of interest. Correlations between regions of interest were calculated. The arctangent of the resulting Pearson’s correlation coefficients were calculated and pair-wise comparisons were made.

### Representational Similarity Analysis

Neurobiological correlates of generalization may arise as similar/dissimilar patterns of activation. To evaluate this possibility, we performed a representational similarity analysis (RSA) using the bilateral CA1 subfield of the hippocampus and dorsal/ventral mPFC as our anatomical regions of interest. The original Test phase model was re-run however this time using un-spatially smoothed functional data – as fine differences in activation patterns were the goal of the RSA analysis.

Representational similarity (voxel-wise correlations) was separately calculated in the anatomical regions of interest between target hits and lure false alarms across the different valences (negative, neutral, positive). Large values indicated similar patterns of activation and small values related to dissimilar patterns of activation between target hits and lure false alarms. To evaluate the relation between behavioral generalization and neural measures of pattern dissimilarity separate correlations were performed. Bonferroni corrections were used to correct for inflated Type I error due to multiple comparisons.

## SUPPLEMENTAL FIGURES AND FIGURE LEGENDS

**Figure S1.**
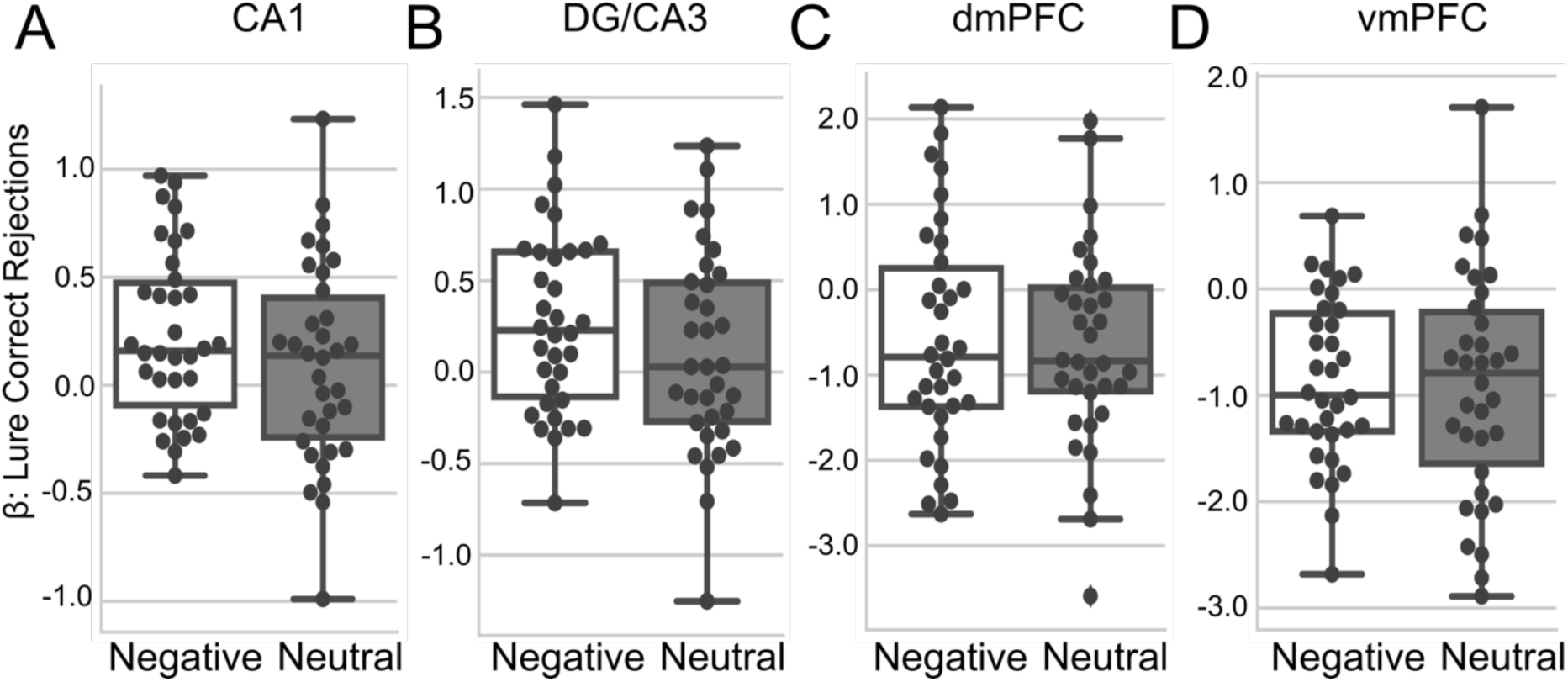
No statistical differences in activation related to discrimination of negative and neutral images across anatomical regions of interest. Anatomical region-of-interest – hippocampal subfield (CA1 and DG/CA3), dorsal medial prefrontal cortex (dmPFC), or ventral medial prefrontal cortex (vmPFC) – examining differences in activations between negative and neutral lure stimuli when they were correctly rejected. (**A**) The CA1 hippocampal subfield (*P* = 0.03) and (**B**) the DG/CA3 subfield (*P* = 0.03) exhibited a trend towards greater activation for negative relative to neutral correct rejections. (**C**) The dmPFC (*P* = 0.76) and (**D**) the vmPFC (*P* = 0.69) did not exhibit significantly different activations between negative and neutral correct rejections. All comparisons were not significant following Bonferroni correction for multiple comparisons.

**Figure S2.**
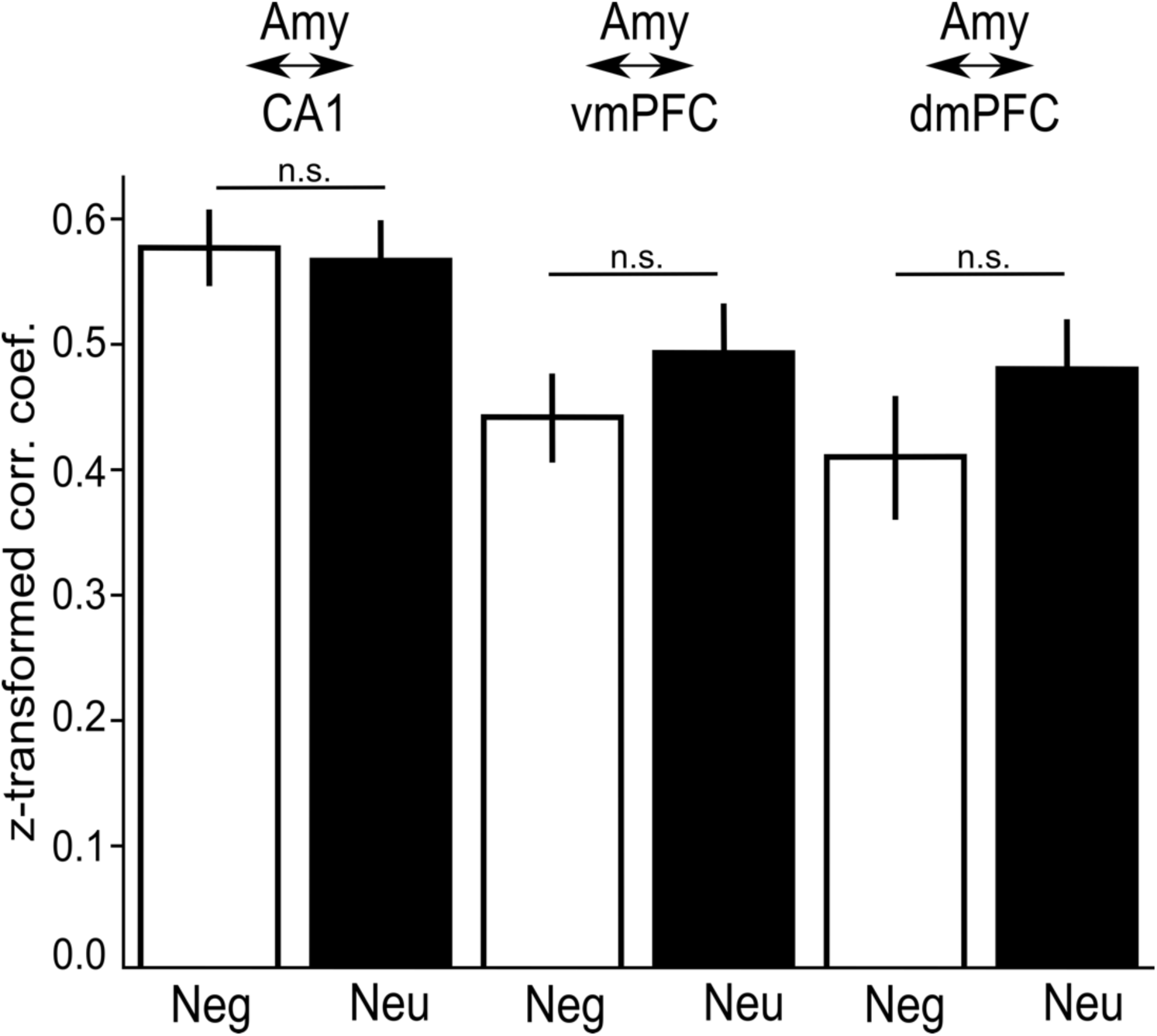
Functional coupling between the amygdala and target regions does not differentiate between negative and neutral scenes when subsequently correctly rejected. Task based functional connectivity during the Study phase for Correctly Rejected negative (Neg) and neutral (Neu) lures between the amygdala (Amy) and three target regions: CA1, ventral medial prefrontal cortex (vmPFC), and dorsal medial prefrontal cortex (dmPFC). Functional coupling for negative and neutral items that were subsequently correctly rejected were not significantly different between any of the a priori anatomically defined regions of interest: Amy ↔ CA1 (*P* = 0.32); Amy ↔ dmPFC (*P* = 0.64); Amy ↔ vmPFC (*P* = 0.45).

**Figure S3.**
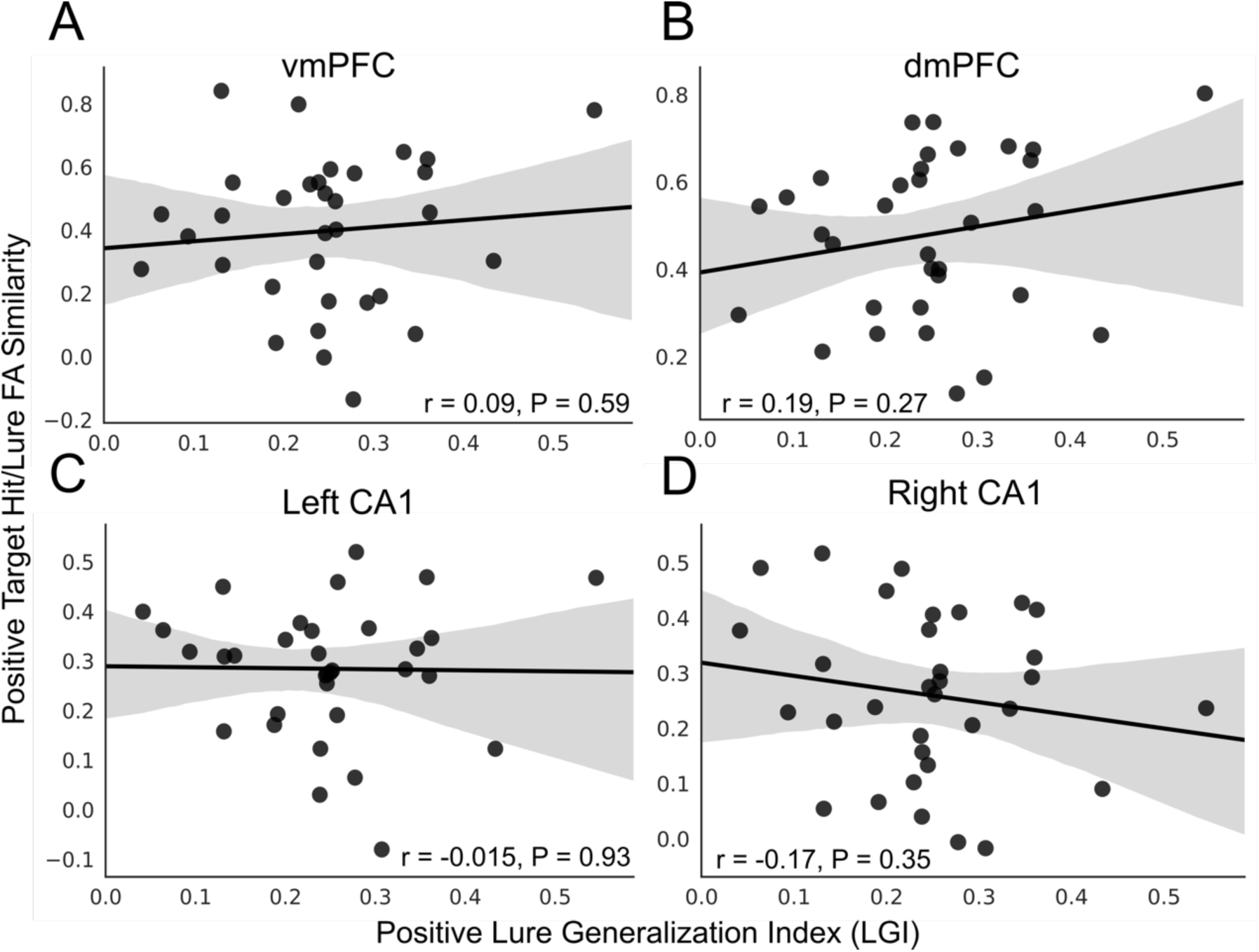
Representational similarity in the mPFC and CA1 not associated with positive overgeneralization. Correlations between positive target hit and lure false alarm (FA) similarity in the (**A**) ventral medial prefrontal cortex (vmPFC), (**B**) dorsal medial prefrontal cortex (dmPFC) and the (**C**) left and (**D**) right CA1 and LGI behavioral performance. No regions exhibited a significant correlation between positive target hit/lure FA similarity and positive LGI performance.

**Figure S4.**
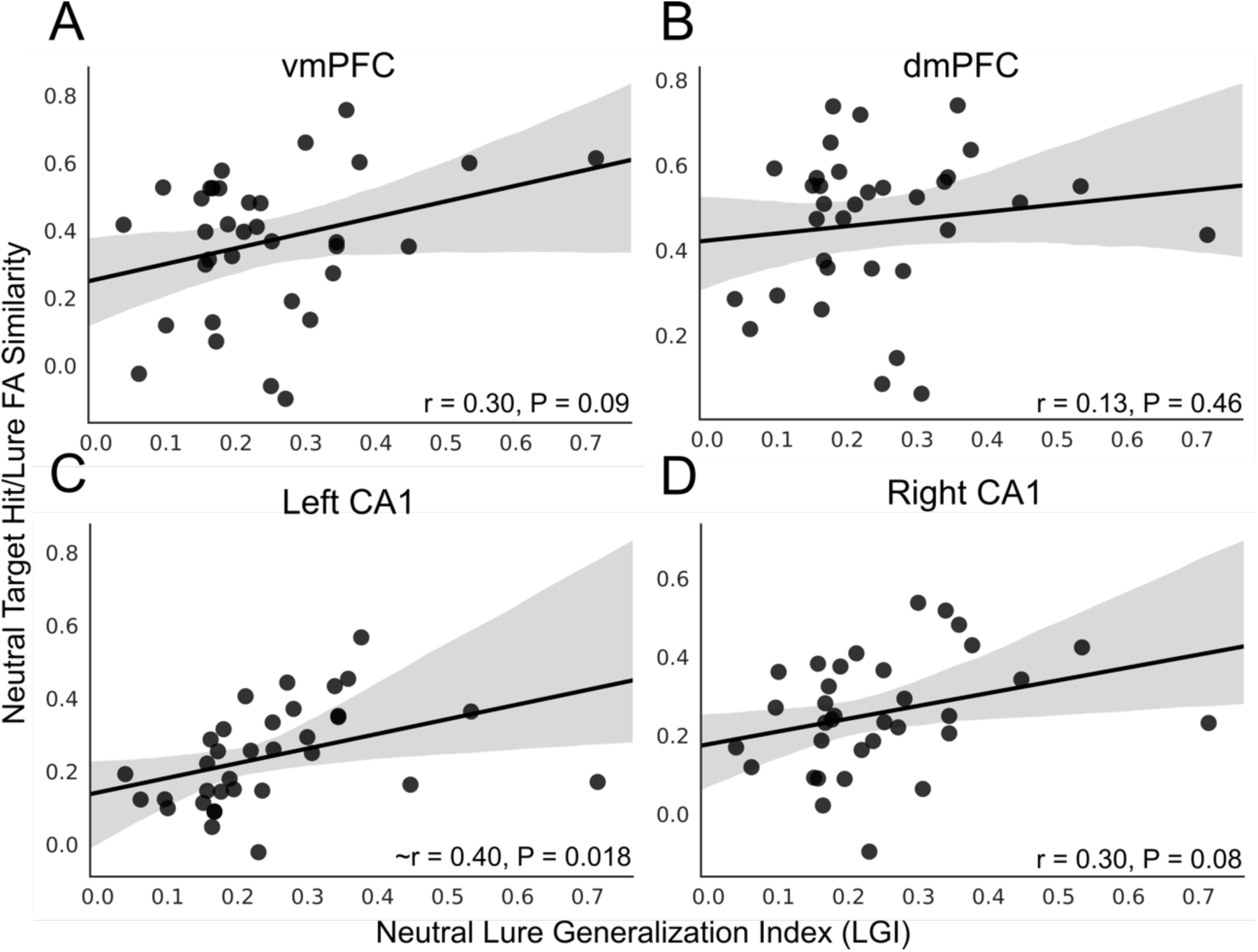
Representational similarity in the mPFC and CA1 not associated with neutral overgeneralization. Correlations between neutral target hit and lure false alarm (FA) similarity in the (**A**) ventral medial prefrontal cortex (vmPFC), (**B**) dorsal medial prefrontal cortex (dmPFC) and the (**C**) left and (**D**) right CA1 and neutral LGI behavioral performance. The vmPFC (*P* = 0.09) and the left (*P* = 0.018) and right (*P* = 0.08) CA1 exhibited a trend in the relation between neutral target hit/lure FA similarity and neutral LGI performance. ∼Trending significant relation.

**Figure S5.**
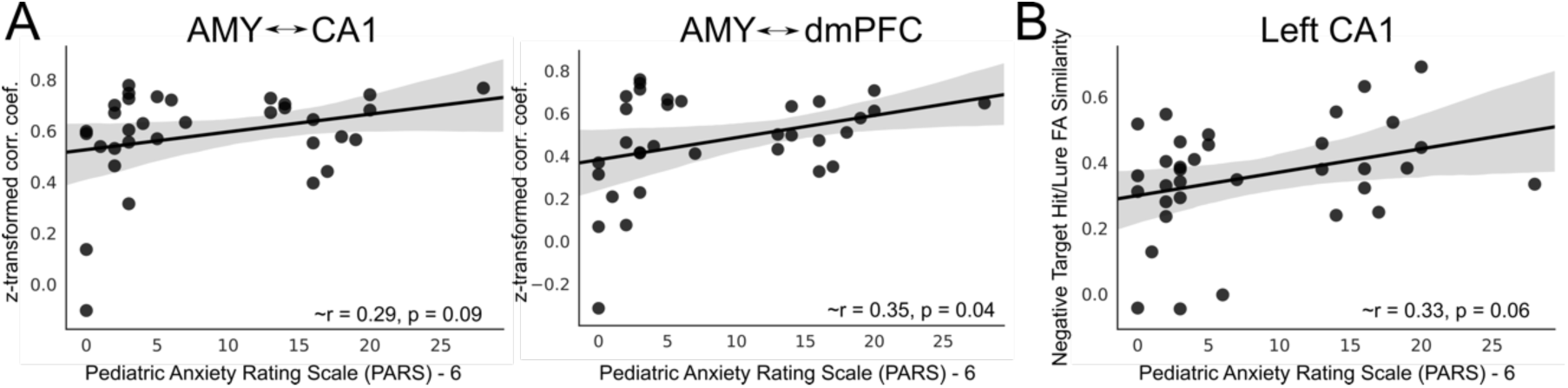
Correlations between select brain measures and anxiety severity as measured by the Pediatric Anxiety Rating Scale (PARS) – 6. (**A**) A trend towards increased functional connectivity between the amygdala and CA1 and the amygdala and dorsal medial prefrontal cortex (dmPFC) during the Study phase for items that were subsequently false alarmed and anxiety severity identified (AMY ↔ CA1: *r* = 0.29, *P* = 0.09; AMY ↔ dmPFC: *r* = 0.35, *P* = 0.04). (**B**) A similar trend was noted in our representational similarity analysis between negative target hits and lure false alarms (FA). More anxious symptoms were related to greater similarity between these trial types in the left CA1 (*r* = 0.33, *P* = 0.06). ∼Trending significant relation.

## Notes

### Competing Interest Statement

The authors have declared no competing interest.

